# Proteomic characterisation of three independent series of sequential cystic fibrosis strains in an International *Pseudomonas aeruginosa* reference panel indicates positive selection in late infection strains

**DOI:** 10.1101/2025.02.14.638290

**Authors:** Joanna Drabinska, Lucia O’ Connor, Caoilin McClean, Siobhán McClean

## Abstract

*Pseudomonas aeruginosa* is a frequent cause of chronic opportunistic infections in people with cystic fibrosis (CF). It is a highly diverse and adaptable Gram-negative bacterium that thrives in many environments. It is well recognised that *P. aeruginosa* adapts over time of colonisation to facilitate chronic infection including loss of virulence factors, however proteomic analyses of the adaptation to chronic infection have been limited. We previously collated and characterised an international panel of *P. aeruginosa* strains from different clinical presentations and geographical regions. Within this panel were three series of sequential isolates from people with CF, comprising eight strains in total which present an excellent opportunity to seek potential conserved adaptations that may be involved in driving chronic colonisation in the CF lung. We compared the proteomes of all eight strains (early versus respective late) to examine whether there were any changes in the proteomes over time of colonisation that were common to the three series of chronic *P. aeruginosa* infection isolates from three different patients. We identified 11 proteins that showed increased abundance in late isolates from all three patient series, many of which are reported to be involved in virulence, regulation of virulence or response to hypoxia and include cystic fibrosis inhibitory factor repressor (CifR), WspR, two two-component response regulators (PA2572 and PA3702), and transcriptional regulator (PA2551). Moreover, we identified three proteins (PA2572, PA3819 and PA5028) that showed increased abundance in all five late isolates from three people with CF. The probability of this being a random event is 5.06 x 10^-53^ and consequently is very strong evidence of positive selection. The increased abundance of PA2573 and PA3819 appears to improve the fitness *P. aeruginosa* to antibiotics and oxidative stress. Overall, despite the diversity within this species, there appear to be common mechanisms of adaptation in the cystic fibrosis lung.

## Introduction

*Pseudomonas aeruginosa* is a Gram-negative bacterium that is a frequent cause of nosocomial and opportunistic infections and that thrives in a wide range of environments, including man-made water systems, contact lens solutions and soil ^1, 2^ . The World Health Organisation have listed *P. aeruginosa* as a high priority pathogen for which new antibiotics are currently needed^3^. It also causes chronic infections in people with cystic fibrosis (CF). Indeed, despite CF patients receiving more than 12 months of modulator therapy, *P. aeruginosa* remains problematic in a substantial cohort of patients and persistent airway inflammation remains ^4,5^. It is clear that chronic infections are not being completely cleared and are more challenging to monitor when patients are on CFTR modulators ^6, 7^

It is well recognised that *P. aeruginosa* adapts over time of infection to facilitate chronic colonisation^1, 8–11^. Some common features of *P. aeruginosa* adaptation reported include loss of virulence factors, including flagella, motility, type III secretion system as recently reviewed ^1^. Given the diversity of *P. aeruginosa* globally, we previously collated an international panel of *P. aeruginosa* 41 strains which had been sequenced ^12^. We subsequently performed a detailed phenotypic analysis and comparative genomic analysis based on whole genome sequence data to define the characteristics of this international panel of strains ^13, 14^. Within this panel, were three independent series of sequential strains that were isolated from three people with CF in geographically different regions. One series of three sequential strains were acquired in Germany from an individual with CF and are designated AA2 (early), AA43 and AA44 (strains 9, 10 and 11 in the panel), with the latter two strains being the last isolates before the patient’s death^15–17^. One was reported as being mucoid (AA43), while AA44 was reported as non-mucoid, however analysis of the panel strains indicated that neither produced alginate on PIA plates and that reversion had occurred^14^. Another series of three strains, designated AMT 0060-1, AMT 0060-2 and AMT 0060-3 were strains from a paediatric patient in Seattle ^18^, with AMT 0060-3 being the early strain when the patient was 7.7 years old and the other two strains being isolated 7.9 years later. Finally a pair of strains (AMT0023-30 and AMT0023-34) were isolated 96 months apart from a young paediatric patient, the first strain was isolated when the patient was only 6 months old^18^. The 96-month strain had 68 unique mutations relative to the first strain. In each case, the time between isolation of the early and late strains was comparable^18, 19^, at approximately 7 years (Table 1).

**Table 1:** Summary of sequential *P. aeruginosa* isolates from three CF patients used in this study and the changes in protein abundance between early and late isolates.

| Series / strain | Source | MLST<sup>13</sup> | Age at first isolation (year) | Time from early strain (year) | Ref | Phenotype characterisation<sup>14</sup> |  |  |  |  |  | # proteins showing changed abundance |
| --- | --- | --- | --- | --- | --- | --- | --- | --- | --- | --- | --- | --- |
|  |  |  |  |  |  | Virulence | Pyocyanin | Swarming | Swimming | Twitching | alginate |  |
| **AA2** | Germany | 708 | 3.9 | - | <sup>17</sup> | - | - | - | - | - | - | - |
| AA43 |  |  |  | 7.5 |  | ↓ | ↓ | ↓ | NC | NC | NC | 72 |
| AA44 |  |  |  | 7.5 |  | ↓ | ↓ | ↓ | NC | NC | NC | 248 |
| **AMT0023-30** | USA | 1394 | 0.5 | - | <sup>18</sup> | - | - | - | - | - | - | - |
| AMT0023-34 |  | 1394 |  | 8 | <sup>18</sup> | ↓ | ↓ | ↓ | Abs | Abs | NC | 182 |
| **AMT0060-3** | USA | 111 | 7.7 | - | <sup>18</sup> | - | - | - | - | - | - | - |
| AMT0060-1 |  | 111 |  | 7.9 |  | ↓ | ↓ | ↓ | ↓ | NC | NC | 166 |
| AMT0060-2 |  | 111 |  | 7.9 |  | ↓ | ↓ | Abs | ↓ | Abs | ↑ | 138 |

The diversity of *P. aeruginosa* is well documented, not only between different environments, but also between different CF patients and indeed within individual CF patients^1, 8, 9, 20–23^. Extensive phenotype characterisation was performed on these sequential CF strains as part of the overall characterisation of the international strain panel^14^. Reduced virulence in the *Galleria mellonella* acute infection model was the only consistently altered phenotype over time of colonisation in all late isolates, in line with other reports ^16, 24, 25^ while reduced pyocyanin was observed in four out of five late strains relative to the earlier strain, with AMT0060-1 being the exception (Table 1). Consequently, this well characterised strain panel presented an excellent opportunity to seek potential conserved alterations in the proteome that may be involved in driving chronic colonisation in the CF lung. Thus, we compared the proteomes of all eight strains to examine whether there were any adaptations that might be common to chronic *P. aeruginosa* and highlight any conserved evolutionary mechanism(s).

## Results and Discussion

### Eleven proteins were changed in abundance in late strains of all CF sequential series highlighting positive selection during chronic infection

Proteomic analysis of each series identified considerable changes in protein abundance between early and late-stage strains (Supplemental Table 1). A total of 138 proteins that had significantly changed abundance by 1.5-fold or more between the AMT0060-2 late strain and the AMT0060-3 early strain, while 166 proteins had changed between the AMT0060-1 late strain and the AMT 0060-3 strain (Table 1). Of these, 57 proteins were common to both late strains, indicating a degree of shared adaptation in this series and as would be expected for strains with a shared ancestor. In the case of the AA2 series a total of 79 proteins were altered in abundance from the AA2 strain to the AA43 strain, while 268 proteins were altered by 1.5-fold or more between AA2 and AA44. Of these 268 proteins, there were only 30 proteins that showed changes in abundance in both late strains in this series. Although only 79 proteins showed altered abundance from AA2 and AA43, 38 % of these changes were maintained in the other late strain AA44. Finally, a total of 182 proteins were altered in abundance by 1.5-fold between AMT-0023-30 and AMT0023-34. Given the diversity across *P. aeruginosa* we specifically sought proteins that were significantly changed in abundance in two or more series of strains to identify possible adaptations that might be conserved during host adaptation within the species.

Detailed analysis of the eight proteomes of the three series of strains indicated that there were only 16 proteins in total that were identified as being altered in abundance in all three series (Figure 1A). Eleven of these were consistently changed across the three series with all showing increased abundance in all later strains of each series (Figure 1B). Consequently, we focussed on the eleven proteins showing increased abundance in all three series (Table 2). These include proteins involved in cell adhesion, cell division, two two-component response regulators (PA2572 and PA3702) and a transcriptional regulator (PA2551). In all cases, (with the sole exception of PA2883, DUF 2897 family protein) these proteins were undetectable in the early strain indicating that the genes were switched on over time of colonisation of the lung (Table 2). Furthermore, in the case of PA2883, while it was detected in AA2, it was undetectable in the other two early strains and is likely that this gene is switched on in the late isolates in these series also. Overall, this indicates a consistent switching on of all 11 of these genes over time of infection. Of the remaining five proteins common to all series, but not showing consistent changes, both the AA2 series and AMT0023 series both showed a decrease in abundance in the later strains, while the AMT0060 series showed increased abundance of all five proteins.

**Figure 1:**
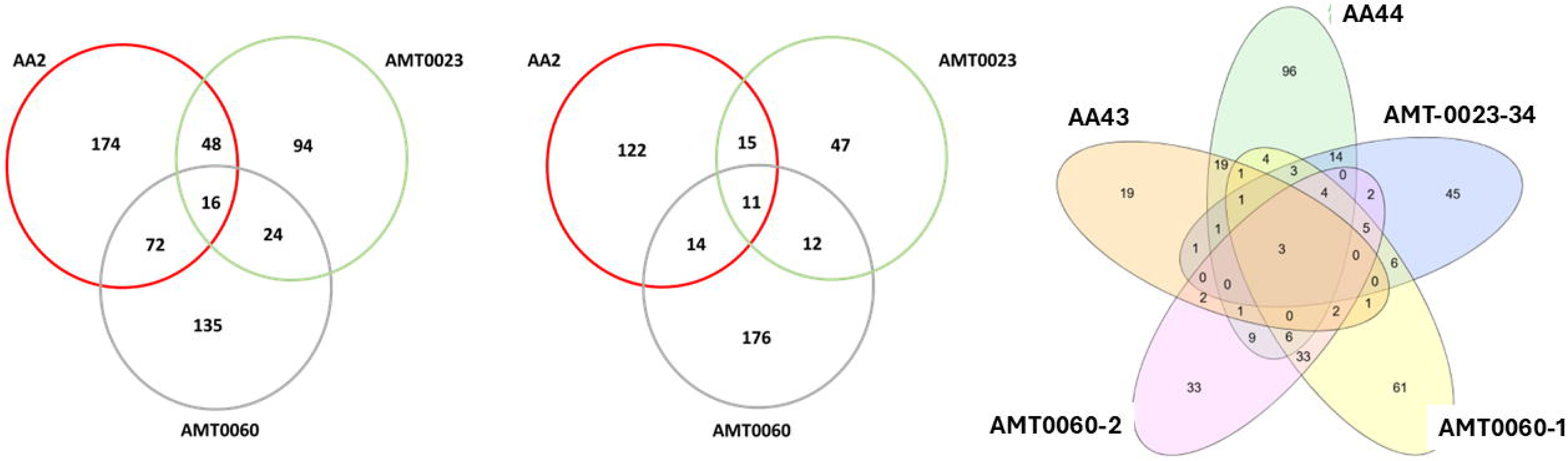
Venn diagram indicting the changes in protein abundance in each independent series of P. aeruginosa sequential strains. Numbers represent the number of proteins in the late isolate(s) relative to the respective early strain in each series. (A) All proteins changed in abundance (i.e. increased or decreased) by 1.5-fold or more in any series of isolates (p < 0.05). (B) The number of proteins that showed increased in abundance by 1.5-fold or more between early and late isolate(s) in any given series (p < 0.05). (C) The numbers of proteins that show increased abundance in any of the five late isolates (p < 0.05).

**Table 2.** Proteins showing increased abundance in later strains in all three series of sequential *P. aeruginosa* strains.

| Gene ID | Protein names | GO (bioprocess/ cellular location) | AA2 | AA43 | AA44 | AMT2<sub>3</sub> | AMT2<sub>3</sub> | AMT0<sub>060-2</sub> | AMT0<sub>060-1</sub> | AMT0<sub>060-3</sub> |
| --- | --- | --- | --- | --- | --- | --- | --- | --- | --- | --- |
| PA1462 | Probable plasmid partitioning protein | Cell division | 0 | 0 | 3 | 0 | 3.7 | 0 | 11 | 11 |
| PA2551 | Probable transcriptional regulator | regulation of DNA-templated transcription | 0 | 0 | 11 | 0 | 15 | 0 | 3.5 | 0 |
| **PA2572** | **Probable two-component response regulator** | phosphorelay signal transduction system; | 0 | 11 | 15 | 0 | 12 | 0 | 18 | 12 |
| PA2679 | Methyltransferase type 11 domain-containing protein | methyltransferase activity | 0 | 16 | 17 | 1/4 | 3.2 | 0 | 16 | - |
| PA2883 | DUF2897 family protein | membrane | 0 | 0 | 2.2 | 1/4 | 3.6 | 0 | 18 | 16 |
| PA2931 | CifR | DNA binding; negative regulation of transcription | 0 | 0 | 10 | 0 | 11 | 0 | 10 |  |
| PA3084 | DUF2797 domain-containing protein | Hypothetical, unclassified | 0 | 0 | 15 | 0 | 15 | 0 | 10 | 13 |
| PA3271 | histidine kinase (EC 2.7.13.3) | phosphorelay signal transduction system | 0 | 0 | 16 | 0 | 13 | 0 | 3.4 |  |
| PA3702 | Probable two-component response regulator | cell adhesion involved in single-species biofilm formation | 0 | 0 | 12 | 0 | 16 | 0 | 15 | 14 |
| **PA3819** | **Glycine zipper 2TM domain-containing protein** | Outer membrane lipoprotein protein SlyB | 0 | 12 | 19 | 0 | 12 | 1/4 | 4 | 5 |
| **PA5028** | **AAA domain-containing protein** | Hypothetical; ParAB_family; Nucleoside triphosphate hydrolase | 0 | 15 | 17 | 0 | 13.6 | 0 | 17 | 13 |
| Not detected |  | Numbers within cells indicate Log<sub>2</sub>-fold change |  |  |  |  |  |  |  |  |
| < 3 |  |  |  |  |  |  |  |  |  |  |
| < 8 |  |  |  |  |  |  |  |  |  |  |
| < 12 |  |  |  |  |  |  |  |  |  |  |
| < 20 |  |  |  |  |  |  |  |  |  |  |

**Table 3.**
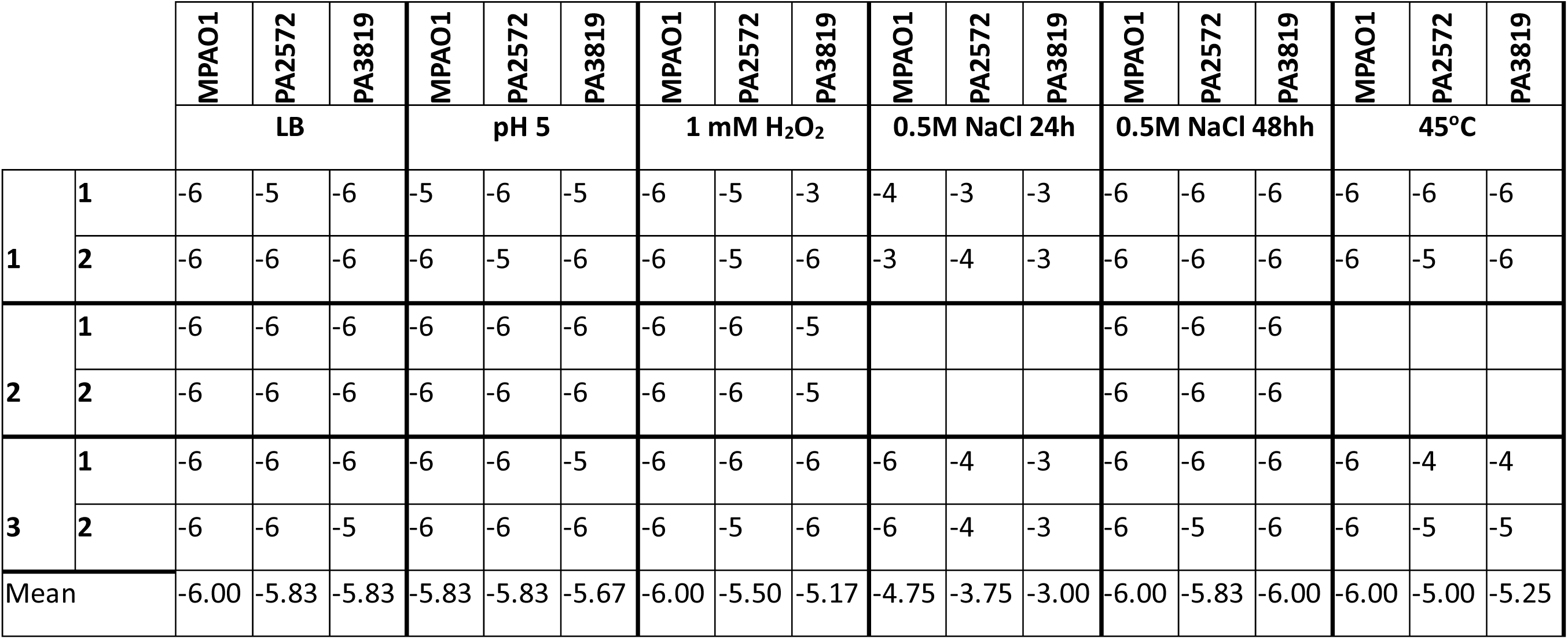
Summary of stress test analysis highlighting the lowest dilutions (10^-x^) at which colonies were visible in three independent experiments.

The AA2 and AMT0023 series shared an additional 48 proteins that were changed in abundance, 42 of which showed consistent change (15 were increased in abundance while 27 were reduced in abundance). A total of 24 additional proteins were common between the two series of strains from the two US paediatric patients (AMT0023 and AMT0060) (Figure 1A). Given that there were 182 proteins showing changes in abundance in the AMT0023 series, 48% of these were also found in one of the other two sets of sequential strains which strongly indicates that despite the diversity inherent in *P. aeruginosa* certain adaptation pathways are conserved over time of colonisation. Moreover, a total of 72 proteins were common between the AA2 series and the AMT0060 series, with 40 of these showing consistent changes, in terms of consistently showing increased or decreased abundance in the later strain. The other 32 proteins showed opposing changes in abundance (increased in one series of sequential strains and decreased in the other).

Overall, 144 proteins showed altered abundance in more than one series of strains (Figure 1). The consistent change in abundance of the same 11 proteins in sequential strains from three different patients suggests that these proteins have roles in the adaptation of *P. aeruginosa* to chronically colonise the CF lung. Of these 11, there were only three proteins which showed increased abundance in all five late isolates from the three series (Figure 1C and table 2). The probability that the same three proteins would show increased abundance in all five late strains from three individual early strains is a random event is 5.06 x 10^-53^ and consequently is strong evidence of positive selection.

### Many proteins that showed increased abundance in late isolates have roles in regulation of virulence factors, response to hypoxia or motility

One of the proteins that shows increased abundance in all five late strains, **PA3819,** is an outer membrane lipoprotein with a glycine zipper domain^26^. It was reported to be a putative SlyB-like lipoprotein that is encoded in the AlgU (σ^22^) regulon and may have a role in membrane integrity ^27^. AlgU is a sigma factor that regulates expression of alginate and is released by cell-wall stress. PA3819 is one of eight lipoproteins activated in mucoid *P. aeruginosa*, many of which are involved in TLR signalling, in late strains ^28^. Given the variability in IL-6 and IL-8 secretion across the five isolates and the previous identification of alginate production in AMT0060-2 only^14^, AlgU upregulation of PA3819 appears to be quite specific, rather than regulon-wide.

**PA2572** is also increased in abundance in all five late strains. It is a HD-GYP domain protein, characterised as a probable two component regulator encoded beside a chemotaxis transducer gene and is predicted to be involved in the regulation of cell motility^26^. It binds directly with the sensor, PA2573, which is co-expressed with PA2572 under several conditions including aerobic growth in the presence of nitrate and in bacteria co-cultured with respiratory epithelia cells. Recent reports suggest that PA2572 and PA2573 are responsible for ExoS production and pyocyanin production^29^. It is described as an orphan chemotaxis regulator which has a role in virulence. Mutants of PA2572 or PA2573 showed considerably reduced virulence in a *G. mellonella* model which contrasts with the observations in the three isolate series, suggesting additional levels of regulation. PA2572 is one of three HD-GYP proteins in *P. aeruginosa*, but unlike the other two it does not hydrolyse c-di-GMP and has a negative influence on swarming. Consistent with this, swarming in BSM-G agar was previously reported to be decreased in all five late strains^14^

**PA5028** and **PA1462** are both cytoplasmic membrane proteins ^26^ that were both switched on in later strains relative to early strains in all three series, with PA5028 showing increased abundance in all five late strains. PA1462 was undetectable in AA43 but showed increased abundance in AA44 and all other late strains. It is predicted to be a cytoplasmic membrane protein that is involved in cell division^26^ and shares homology with the ATPase ParA^30^. In *Vibrio cholerae* ParA has been associated with chemotaxis. PA5028 is predicted to be a P-loop containing nucleoside triphosphate hydrolase with an AAA domain^26^. It is also classified as a ParAB family protein. Interestingly both PA1462 and PA2058 have been identified as partners for ParB (PA5562, spoOJ), a DNA binding protein. Par proteins form partitioning systems which contribute not only to cell division, but also are also global regulators of multiple proteins^31,32^.

**WspR (PA3702)** is a diguanylate cyclase which stimulates c-di-GMP production and acts as the response regulator of the Wsp system which showed increased abundance in all late strains except AA43. The Wsp system is a surface sensing system that responds to cell envelope stress. It negatively regulates flagellum-dependent cell motility and promotes biofilm formation^33^. In all series the protein changed from being undetectable in the early strain, again suggesting that the gene was induced in late strains in all three series. The increased abundance of WspR in all series of strains is consistent with the loss of motility widely reported for *P. aeruginosa* chronic isolates^10, 34, 35^.

**CifR** is an TetR-family, epoxide responsive repressor (CFTR inhibitory factor) which showed increased abundance in one late isolate in each of the three series. It regulates the expression of the virulence factor Cif via direct binding and repression^36^. Cif reduces apical expression of ABC transporters including CFTR by enhancing their polyubiquitination and lysosomal degradation. Ballok et al., have previously shown that *P. aeruginosa* CF isolates maintain CifR expression over time ^36^, consistent with our observations in the three independent series of strains. Interestingly, given the diversity across *P. aeruginosa*, the fold change between the early strain and the respective late strain is equivalent, i.e. 10 to 11-fold in all three series.

The probable transcriptional regulator, **PA2551 is** predicted to be a DNA-binding protein which shares homology with LysR family transcriptional regulators and a component of an intricate network of transcriptional regulators. Recent reports suggest that PA2551 inhibits BrsA, a negative regulator of pyruvate metabolism and the TCA cycle. The latter has been reported to be released in response to stress and a requirement to cope with adverse conditions that can manifest in slow growth. It is possible that PA2551 acts to counteract this slowdown and maintain growth in stressful environment of the CF lung.

**PA2679** is a hypothetical protein and predicted to have methyl transferase activity and showed increased abundance in all late strains except AA43. Its expression was reported to be enhanced in response to hypoxia (0.4% oxygen) and moderately enhanced in response to airway epithelia ^37^. Another hypothetical protein **PA2883** is predicted to be a cytoplasmic membrane protein was also increased in all late strains except AA43. STRING analysis suggested co-expression with PA4608 (methyltransferase-associated PilZ, MapZ), a c-diGMP binding protein^38^. The GEO expression database indicates that its expression doubled in response to hypoxia and increased dramatically in response to airway epithelia^37^ . *In vivo* transcriptome analysis showed it is induced during infection and in response to antimicrobials. PA3084 is another hypothetical protein that was identified as a conditional essential gene for cardiomyocyte infection in the absence of amikacin ^39^. While all three hypothetical proteins were identified in all three series, neither PA2679, PA2883, nor PA3084 were identified in the AA43 strain highlighting a distinct evolutionary path for this late strain.

**PA3271** is classified as a probable two component sensor with kinase activity and is predicted to be a cytoplasmic membrane protein ^26^. A **PA3271** transposon mutant showed the upregulation of 15 genes and downregulation of 40 genes relative to the wild-type strain^40^. Among the downregulated genes were genes involved in virulence, i.e. PchF (pyochelin synthase) and PchD (pyochelin biosynthesis protein); elastase, PvdS, FliD, flagellar capping protein, TonB receptor, suggesting PA3271 is likely to regulate virulence. Consistent with that report, the abundance of PchF, elastase, PvdS, TonB receptor and two flagellar proteins FliL and FliD were also altered in abundance in the late CF strains (Supplemental Table 1).

In summary, the 11 proteins that were consistently increased in abundance in the late isolates from all three series have roles in virulence or regulation of virulence factors, regulation of motility or responses to hypoxia. While eight of the 11 proteins were not identified in all five late strains, the probability of all these eight proteins showing increased abundance in at least one late strain in each patient being random is 2.92 x 10^-9^ in a genome of 5570 genes^41^. Thus, the evolving population may have relied on the expression of these eight proteins in a model of cheating or cooperation and thus the expression of these proteins in three independent sequential series strongly indicates that they are also important for persistence and chronicity. It is interesting that no antibiotic resistance mechanisms are represented among the 11 proteins, despite multiple antibiotic resistance proteins showing increased abundance in all late strains. For example, MexA only increased in the AA2 series, with dramatic increases in abundance in strains AA43 and AA44, but was not detected in the AMT 0060 series and was reduced in abundance in AMT23-0034. The lack of consistent change in antibiotic resistance proteins may be due in part to different antibiotic regimens used.

In addition to these shared proteins, comparison of the biological processes in all three series that were associated with proteins of either increased or decreased in abundance were remarkably similar (Figure 2). In all cases, PANTHER could not assign a biological process to between a third to half of the proteins; however, there were a comparable proportion of proteins in cellular processes, biological regulation processes and metabolic process increased as they were decreased in all three series. While the proportion of proteins associated with localisation was greater in the later strains from the AA and AMT0023 series, there were no proteins associated with localisation in the later strains of the AMT0060 series. There were proteins associated with homeostatic processes showing increased abundance in the AMT0060 and AA2 series but not the AMT0023 series. Similarly, no clear metabolic function appeared to consistently show increased or decreased abundance all three series (Fig 2B). Proteins associated with transporter activity seemed to be more represented among proteins showing reduced abundance in AMT0023 and AA2 late strains, but this was not apparent in the AMT0060 series. Comparison of protein class did not show any consistent pattern of increased or decreased abundance across the three series. This again suggests that the specific 11 proteins that show increased abundance in all three series have an important role in the adaption to chronic infection.

**Figure 2:**
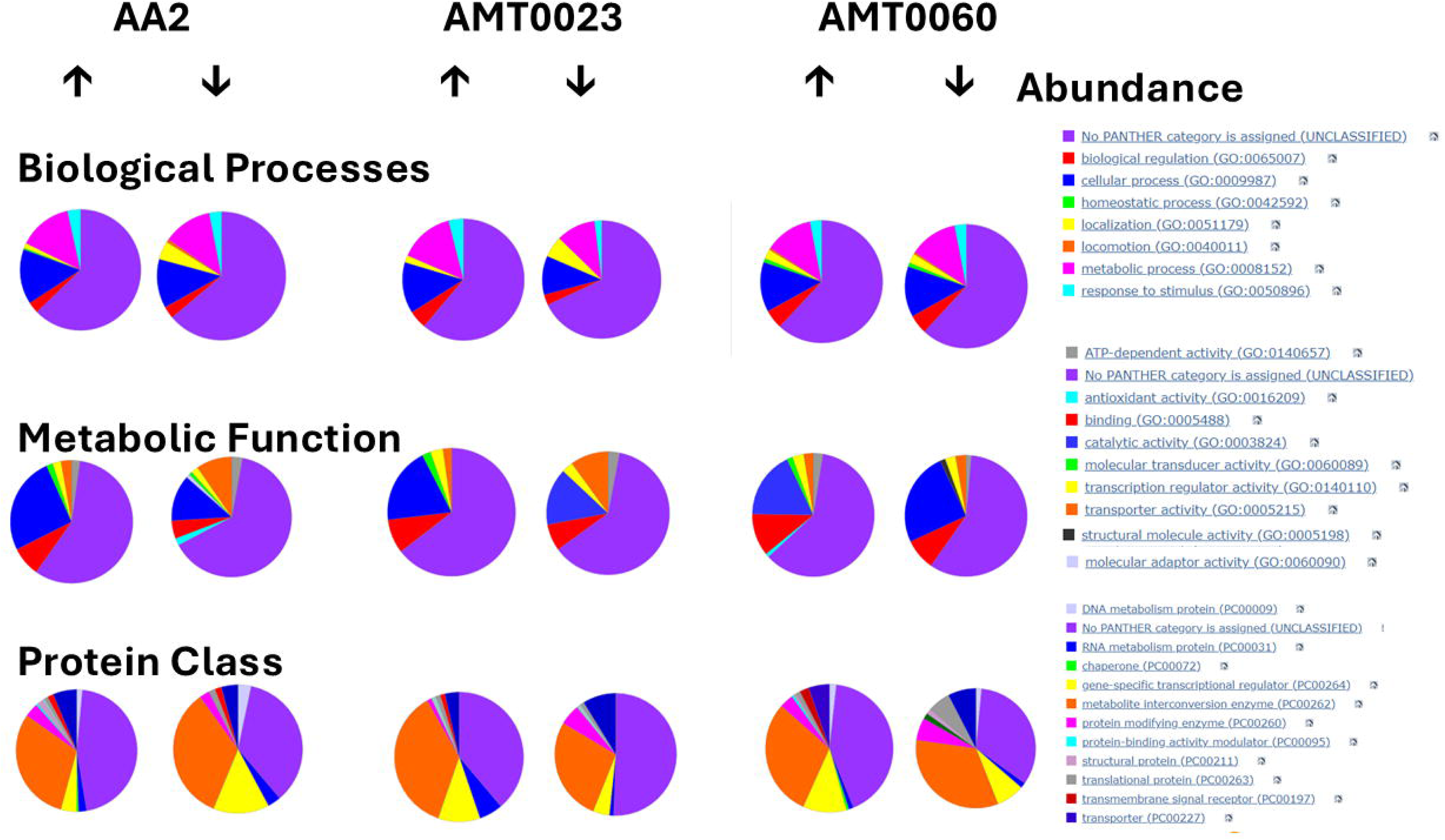
PANTHER categories of proteins that are changed in abundance in each series of sequential P. aeruginosa CF strains. Up and down arrows indicate increased or decreased abundance, respectively. Biological processes, metabolic functions and protein classes are denoted in the respective colour-coded keys.

Previous studies comparing sequential isolates have not identified these 11 proteins as being consistently increased in abundance. This may be in part due to the use of less sensitive comparative proteomic methods such as 2D electrophoresis in the comparison of sequential CF isolates ^42, 43^ . Indeed, the proteomic analysis of sequential *P. aeruginosa* isolates has been quite limited.

### PA2572 and PA3819 may confer increased fitness in response to stress and reduced antibiotic susceptibility

The increased abundance of PA2572, PA3819 and PA5028 in all five late strains relative to the three respective earlier strains pointed to these proteins being critical to chronic infection. Deletion mutants for PA2572 and PA3819 were available ^44^ and neither were identified as essential genes^45^. Therefore, to probe the potential roles of these two proteins further, we examined the effect of stress responses, including those that are considered typical in the CF lung, in the ΔPA2572 or ΔPA3819 gene deletion mutants to investigate whether increased abundance of either protein might enhance fitness in these conditions. Survival of both transposon mutants was compared with that of MPAO1 under an array of stress conditions. In the absence of stress, survival was comparable between the control strain MPAO1 and the two mutants PA2572 and PA3819 (Figure 3). The ΔPA3819 mutant showed impaired survival in 1mM H_2_O_2_ while the ΔPA2572 mutant was less affected (Figure 3). Both strains also showed impaired survival in osmotic stress conditions (0.5M NaCl) and elevated temperature (45°C) relative to the control. Survival of both mutant strains was unchanged in the presence of mild acid (pH5) relative to MPAO1. The findings that the absence of PA3819 impacts on fitness to survive in oxidative stress conditions, while the absence of both PA3819 and PA2572 appears to impair survival in elevated salt and strongly suggests that the increased abundance of these proteins is likely to support the ability of *P. aeruginosa* to thrive during chronic infection.

**Figure 3:**
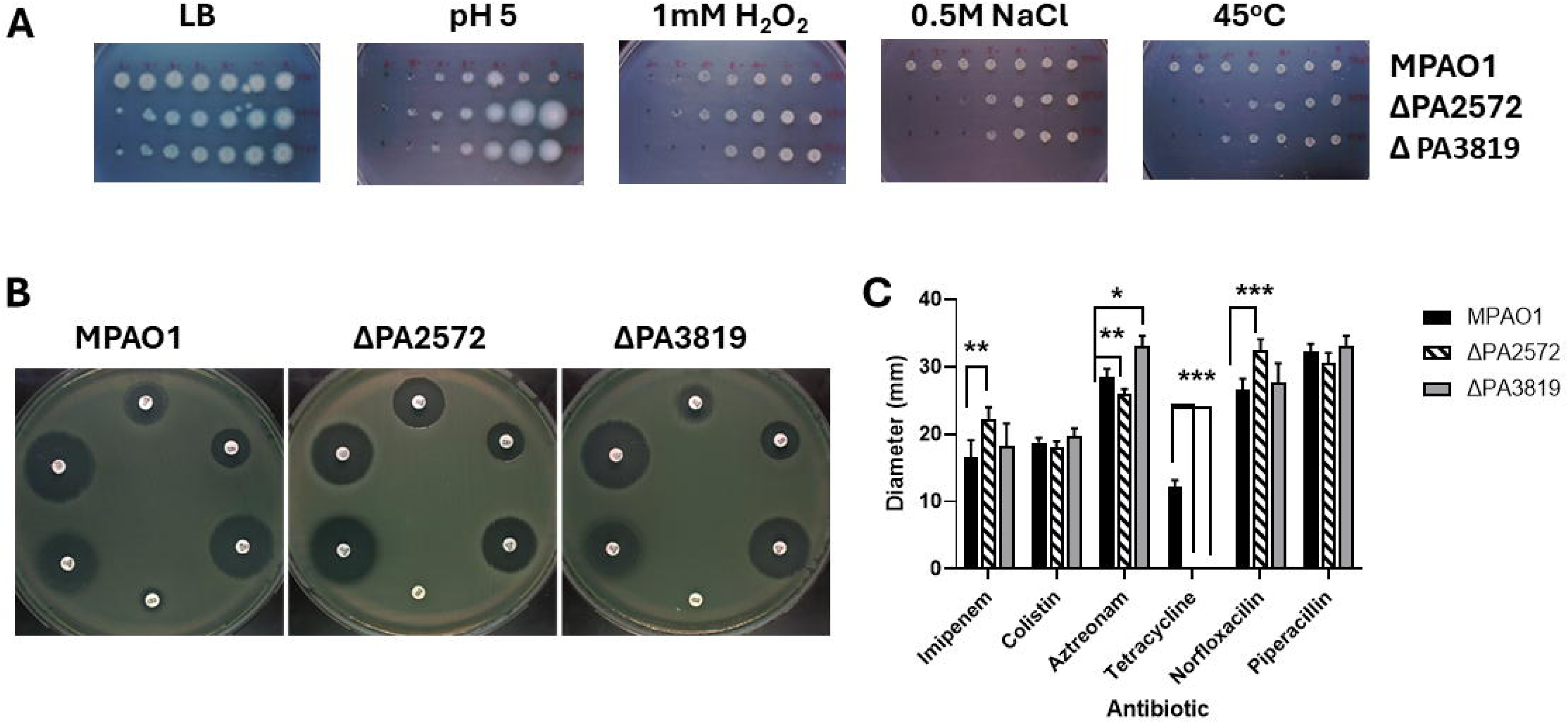
Comparison of ΔPA2572 and ΔPA3819 mutant strains with wild type strain MPAO1 in response to stress or antibiotics. **A)** Survival of ΔPA2572 and ΔPA3819 mutant strains in response to acidic, osmotic, temperature or oxidative stress. **B and C)** Antibiotic susceptibility of ΔPA2572 and ΔPA3819 mutant strains relative to MPAO1 control. **B)** Representative images of plates showing changes in zones of clearance in response to a range of antibiotics, clockwise from top: imipenem, colistin, aztreonam, tetracycline, norfloxacin, piperacillin. **C)** Diameters of zones of clearance in MPAO1, ΔPA2572 and ΔPA3819 mutant strains in response to antibiotics.

Finally, while no antibiotic resistance-associated proteins were consistently changed in all five late strains, we examined whether these two genes confer any fitness in the context of reduced susceptibility to antibiotics commonly used in treatment of CF patients. Increased susceptibility to imipenem, aztreonam and norfloxacin was apparent in the ΔPA2573 mutant while only increased susceptibility to Aztreonam was observed in the ΔPA3819 mutant (Figure 3B and 3C). Interestingly, both mutants showed were resistant to tetracycline. Collectively, this indicates that the increased abundance of these two proteins, and particularly PA2573, in the late isolates may contribute directly or indirectly to the enhanced antibiotic resistance these late strains.

## Conclusion

Given the diversity between *P. aeruginosa* strains and isolates, not only between hosts, but also within an individual host, the increased abundance of eleven proteins in the late strains of all three independent series of sequential strains is clear evidence of positive selection of these proteins. It is likely that there are complex mechanisms at play in the adaptation to chronic infection and undoubtedly there may also be interplay between these proteins and pathways. Thus, this investigation of the individual proteins only provides a snapshot of the overall dynamic adaptation process. Nevertheless, the increased abundance of PA2573 and PA3819 appears to improve the fitness *P. aeruginosa* to antibiotics and oxidative stress, respectively, at least in part. Each of the 11 proteins identified represent targets for future adjuvant therapy to impair the development of chronic infection. Consequently, detailed evaluation of the roles of these proteins in the adaptation process is warranted.

## Materials and Methods

### Bacterial strains

The bacterial strains used in this study are well-characterised strains in the International *P. aeruginosa* panel ^12^ (Table 1). All strains were routinely grown in LB broth (Sigma-Aldrich, St. Louis, MO, USA) at 37°C with orbital agitation (200 rpm) or plated on LB agar (Sigma-Aldrich, St. Louis, MO, USA) and grown at 37°C.

### Whole cell proteome analysis

Whole cell lysates were extracted from cultures by resuspending in 1 ml of ice-cold lysis buffer containing 40mM Tris-HCl (pH 7.8) and 1x cOmplete™, EDTA-free protease inhibitor cocktail (Roche, Basel, Switzerland). Cells were disrupted by a sonication with a sonic dismembrator and microtip 6mm probe (model 505; Fisherbrand, Thermo Fisher Scientific, Waltham, MA, USA) (set with 20% output power using eight 10 s bursts on ice). Cell debris and any remaining intact cells were pelleted using centrifugation (12000 g for 30 min). Supernatants were transferred into fresh tubes and the proteins were precipitated with four volumes of methanol and incubated in -80°C overnight. The samples were centrifuged at 12000 g for 30 min at 4°C, supernatants were removed, and the protein pellets were resuspended in 200 μl of resuspension buffer containing 8 M urea, 50 mM Tris-HCL (pH 8.0), before 1 M 1,4-dithiothreitol (DTT) was added to each supernatant (10 μl/ml lysate) and incubated at 56°C for 30 min, followed by the addition of 1 M iodoacetamide (55 μl/ml lysate) which was incubated in darkness at room temperature for 20 min. Lysates were dialyzed in SnakeSkin™ tubing (Thermo Fisher Scientific, Waltham, MA, USA) with a cut-off of 3.5 kDa against 100 mM ammonium bicarbonate overnight, with stirring at 4°C, followed by at least 4 more hours incubation with fresh volumes of ammonium bicarbonate. Trypsin (400 ng/μl) (Sigma-Aldrich, St. Louis, MO, USA) was added to the dialyzed protein (5 μl /100 μl protein) and tubes were incubated overnight at 37°C. Aliquots from each sample were placed in fresh tubes and samples were dried in a vacufuge concentrator system (model 5301; Eppendorf, Hamburg, Germany) at 65°C and resuspended in 20 μl of ZipTips resuspension buffer containing 0.5% trifluoroacetic acid (TFA) in MilliQ water. ZipTips (Merc Millipore, Burlington, MA, USA) were used for peptide purification as per the manufacturers’ instructions. Nanodrop spectrophotometer DS-11+ (DeNovix, Wilmington, DE, USA) was used for peptide concentration measurements. The samples were analysed on a Bruker TimsTOF Pro mass spectrometer connected to a Evosep One chromatography system. Peptides were separated on an 8 cm analytical C18 column (Evosep, 3 µm beads, 100 µm ID) using the pre-set 33 samples per day gradient on the Evosep one. A trapped ion mobility (TIMS) analyser was synchronized with a quadrupole mass filter to enable the highly efficient PASEF (Parallel Accumulation Serial Fragmentation acquisition) procedure with acquisition rates of 100 Hz. The accumulation and ramp times for the TIMS were both set to 100 ms., with an ion mobility (1/k0) range from 0.62 to 1.46 Vs/cm. Spectra were recorded in the mass range from 100 to 1,700 m/z. The precursor (MS) Intensity Threshold was set to 2,500 and the precursor Target Intensity set to 20,000. Each PASEF cycle consisted of one MS ramp for precursor detection followed by 10 PASEF MS/MS ramps, with a total cycle time of 1.16 s.

Protein identification and label-free quantitative (LFQ) analysis were conducted using MaxQuant (v 2.4.2.0, https://maxquant.org/) by searching against the reference strain *P. aeruginosa* PAO1 (ATCC 15692). For protein identification, the following search parameters were used: trypsin was the digesting enzyme with up to two missed cleavages allowed for; oxidation on methionine and acetylation on the N terminus were selected as variable modifications with a fixed modification of carbamidylation on cysteine; Label Free Quantitation (LFQ) and Match Between Runs (MBR) options were selected. FDR for protein and peptide were set at 1%.

### Stress sensitivity assays

Overnight cultures of PA strains were diluted to a CFU/ml of 10^8^ and a range of serial dilutions was prepared. Suspensions (1ul) were plated on one of the following: LB agar control plates, LB agar supplemented with 0.5 M NaCl (osmotic stress), LB agar supplemented with 1mM H_2_O_2_ (oxidative stress), or LB agar, pH 5 (acidic stress). The plates were incubated at 37°C for 24 h. Temperature stress was determined by incubation of an LB agar plate at 45°C for 24 h. Change of sensitivity was determined by determining the lowest dilution at which bacterial growth was observed. The experiment was repeated three independent times.

### Determination of antibiotic Resistance

Overnight bacterial cultures were diluted in 10 ml of fresh LB to OD600 equal to 0.1. Bacteria were spread evenly on Mueller–Hinton agar plates (Neogen, Lansing, MI, USA) using cotton swabs, in three directions, to give a homogenous lawn. Antibiotic disks (Oxoid, Hampshire, UK) were placed on the surface of the agar and the plates were incubated overnight in normoxic conditions at 37°C. Growth inhibition zone diameters were measured in at least two points, and the mean was used for statistical analysis. The experiment was repeated three times.

### Statistical and probability analysis

Statistical analysis was performed by One-Way Analysis of Variance (ANOVA) using Prism software with the Kruskal-Wallis test when the data did not pass the Shapiro-Wilk normality test. All graphs and were prepared using GraphPad Prism 8.0.2.

To determine the probability of all 5 late isolates in 3 patients undergoing the same adaptation, the following calculation was performed: (reciprocal of (# genes / 3 patients)) number of isolates

## Acknowledgements.

JJ and LOC were supported by an SFI Frontiers of the Future award (20/FFP-P/8717) to SMcC. The authors are grateful to Ciarán J. Carey for their critical reading of the manuscript and to Dr David Gomez-Matallanas for his advice in proteomic analysis and for the support of the staff and facilities in the UCD Conway proteomics core.

## References

[1] Jurado-Martin, I., Sainz-Mejias, M., and McClean, S. (2021) Pseudomonas aeruginosa: An Audacious Pathogen with an Adaptable Arsenal of Virulence Factors, Int J Mol Sci 22.

[2] Rutherford, V., Yom, K., Ozer, E. A., Pura, O., Hughes, A., Murphy, K. R., Cudzilo, L., Mitchell, D., and Hauser, A. R. (2018) Environmental reservoirs for exoS+ and exoU+ strains of Pseudomonas aeruginosa, Environ Microbiol Rep 10, 485–492.

[3] WHO. (2024) Bacterial Priority Pathogens List, 2024: bacterial pathogens of public health importance to guide research, development and strategies to prevent and control antimicrobial resistance, Licence: CC BY-NC-SA 3.0 IGO. ed., Geneva: World Health Organization.

[4] Armbruster, C. R., Hilliam, Y. K., Zemke, A. C., Atteih, S., Marshall, C. W., Moore, J., Koirala, J., Krainz, L., Gaston, J. R., Lee, S. E., Cooper, V. S., and Bomberger, J. M. (2024) Persistence and evolution of Pseudomonas aeruginosa following initiation of highly effective modulator therapy in cystic fibrosis, mBio 15, e0051924.

[5] Saluzzo, F., Riberi, L., Messore, B., Lore, N. I., Esposito, I., Bignamini, E., and De Rose, V. (2022) CFTR Modulator Therapies: Potential Impact on Airway Infections in Cystic Fibrosis, Cells 11.

[6] Nichols, D. P., Paynter, A. C., Heltshe, S. L., Donaldson, S. H., Frederick, C. A., Freedman, S. D., Gelfond, D., Hoffman, L. R., Kelly, A., Narkewicz, M. R., Pittman, J. E., Ratjen, F., Rosenfeld, M., Sagel, S. D., Schwarzenberg, S. J., Singh, P. K., Solomon, G. M., Stalvey, M. S., Clancy, J. P., Kirby, S., Van Dalfsen, J. M., Kloster, M. H., Rowe, S. M., and group, P. S. (2022) Clinical Effectiveness of Elexacaftor/Tezacaftor/Ivacaftor in People with Cystic Fibrosis: A Clinical Trial, Am J Respir Crit Care Med 205, 529–539.

[7] Allen, L., Allen, L., Carr, S. B., Davies, G., Downey, D., Egan, M., Forton, J. T., Gray, R., Haworth, C., Horsley, A., Smyth, A. R., Southern, K. W., and Davies, J. C. (2023) Future therapies for cystic fibrosis, Nat Commun 14, 693.

[8] Colque, C. A. a. (2020) Hypermutator pseudomonas aeruginosa exploits multiple genetic pathways to develop multidrug resistance during long-term infections in the airways of cystic fibrosis patients, Antimicrobial Agents and Chemotherapy 64.

[9] Winstanley, C., O’Brien, S., and Brockhurst Michael, A. (2016) Pseudomonas aeruginosa Evolutionary Adaptation and Diversification in Cystic Fibrosis Chronic Lung Infections, pp 327–337.

[10] Cullen, L., and McClean, S. (2015) Bacterial Adaptation during Chronic Respiratory Infections, Pathogens 4, 66–89.

[11] Sousa, A. M., and Pereira, M. O. (2014) Pseudomonas aeruginosa Diversification during Infection Development in Cystic Fibrosis Lungs-A Review, Pathogens 3, 680–703.

[12] De Soyza, A., Hall, A. J., Mahenthiralingam, E., Drevinek, P., Kaca, W., Drulis-Kawa, Z., Stoitsova, S. R., Toth, V., Coenye, T., Zlosnik, J. E., Burns, J. L., Sa-Correia, I., De Vos, D., Pirnay, J. P. T. J. K., Reid, D., Manos, J., Klockgether, J., Wiehlmann, L., Tummler, B., McClean, S., Winstanley, C., and On behalf of, E. U. F. P. f. C. A. B. M. C. s. v. d. o. c. f. p. (2013) Developing an international Pseudomonas aeruginosa reference panel, Microbiologyopen.

[13] Freschi, L., Bertelli, C., Jeukens, J., Moore Matthew, P., Kukavica-Ibrulj, I., Emond-Rheault Jean, G., Hamel, J., Fothergill Joanne, L., Tucker Nicholas, P., McClean, S., and Klockgether Jens, a. (2018) Genomic characterisation of an international Pseudomonas aeruginosa reference panel indicates that the two major groups draw upon distinct mobile gene pools, FEMS Microbiology Letters 365.

[14] Cullen, L., Weiser, R., Olszak, T., Maldonado, R. F., Moreira, A. S., Slachmuylders, L., Brackman, G., Paunova-Krasteva, T. S., Zarnowiec, P., Czerwonka, G., Reilly, J., Drevinek, P., Kaca, W., Melter, O., De Soyza, A., Perry, A., Winstanley, C., Stoitsova, S. R., Lavigne, R., Mahenthiralingam, E., Sa-Correia, I., Coenye, T., Drulis-Kawa, Z., Augustyniak, D., Valvano, M. A., and McClean, S. (2015) Phenotypic characterization of an international Pseudomonas aeruginosa reference panel: strains of cystic fibrosis (CF) origin show less in vivo virulence than non-CF strains, Microbiology 161, 1961–1977.

[15] Bragonzi, A., Farulla, I., Paroni, M., Twomey, K. B., Pirone, L., Lore, N. I., Bianconi, I., Dalmastri, C., Ryan, R. P., and Bevivino, A. (2012) Modelling co-infection of the cystic fibrosis lung by Pseudomonas aeruginosa and Burkholderia cenocepacia reveals influences on biofilm formation and host response, PLoS One 7, e52330.

[16] Bragonzi, A., Paroni, M., Nonis, A., Cramer, N., Montanari, S., Rejman, J., Di Serio, C., Doring, G., and Tummler, B. (2009) Pseudomonas aeruginosa microevolution during cystic fibrosis lung infection establishes clones with adapted virulence, Am J Respir Crit Care Med 180, 138–145.

[17] Bragonzi, A., Wiehlmann, L., Klockgether, J., Cramer, N., Worlitzsch, D., Doring, G., and Tummler, B. (2006) Sequence diversity of the mucABD locus in Pseudomonas aeruginosa isolates from patients with cystic fibrosis, Microbiology (Reading) 152, 3261–3269.

[18] Mulcahy, L. R., Burns, J. L., Lory, S., and Lewis, K. (2010) Emergence of Pseudomonas aeruginosa strains producing high levels of persister cells in patients with cystic fibrosis, J Bacteriol 192, 6191–6199.

[19] Tümmler, B. (2019) Emerging therapies against infections with Pseudomonas aeruginosa, F1000Research 8.

[20] Jorth, P., Staudinger Benjamin, J., Wu, X., Hisert Katherine, B., Hayden, H., Garudathri, J., Harding Christopher, L., Radey Matthew, C., Rezayat, A., Bautista, G., Berrington William, R., Goddard Amanda, F., Zheng, C., Angermeyer, A., Brittnacher Mitchell, J., Kitzman, J., Shendure, J., Fligner Corinne, L., Mittler, J., Aitken Moira, L., Manoil, C., Bruce James, E., Yahr Timothy, L., and Singh Pradeep, K. (2015) Regional Isolation Drives Bacterial Diversification within Cystic Fibrosis Lungs, Cell Host and Microbe 18, 307--319.

[21] Darch, S. E., McNally, A., Harrison, F., Corander, J., Barr, H. L., Paszkiewicz, K., Holden, S., Fogarty, A., Crusz, S. A., and Diggle, S. P. (2015) Recombination is a key driver of genomic and phenotypic diversity in a Pseudomonas aeruginosa population during cystic fibrosis infection, Sci Rep 5, 7649.

[22] Feliziani, S., Marvig Rasmus, L., Lujn Adela, M., and Moyano Alejandro, J. a. (2014) Coexistence and Within-Host Evolution of Diversified Lineages of Hypermutable Pseudomonas aeruginosa in Long-term Cystic Fibrosis Infections, PLoS Genetics 10, e1004651.

[23] Workentine, M. L., Sibley, C. D., Glezerson, B., Purighalla, S., Norgaard-Gron, J. C., Parkins, M. D., Rabin, H. R., and Surette, M. G. (2013) Phenotypic heterogeneity of Pseudomonas aeruginosa populations in a cystic fibrosis patient, PLoS One 8, e60225.

[24] Cullen, L., Weiser, R., Olszak, T., Maldonado Rita, F., Moreira Ana, S., Slachmuylders, L., Brackman, G., Paunova-Krasteva Tsvetelina, S., Zarnowiec, P., Czerwonka, G., Reilly, J., Drevinek, P., Kaca, W., and Melter Oto, a. (2015) Phenotypic characterization of an international Pseudomonas aeruginosa reference panel: Strains of cystic fibrosis (CF) origin show less in vivo virulence than non-CF strains, Microbiology (United Kingdom) 161, 1961–1977.

[25] Lore, N. I., Cigana, C., De Fino, I., Riva, C., Juhas, M., Schwager, S., Eberl, L., and Bragonzi, A. (2012) Cystic fibrosis-niche adaptation of Pseudomonas aeruginosa reduces virulence in multiple infection hosts, PLoS One 7, e35648.

[26] Winsor, G. L., Lo, R., Ho Sui, S. J., Ung, K. S., Huang, S., Cheng, D., Ching, W. K., Hancock, R. E., and Brinkman, F. S. (2005) Pseudomonas aeruginosa Genome Database and PseudoCAP: facilitating community-based, continually updated, genome annotation, Nucleic acids research 33, D338–343.

[27] Wood, L. F., and Ohman, D. E. (2009) Use of cell wall stress to characterize sigma 22 (AlgT/U) activation by regulated proteolysis and its regulon in Pseudomonas aeruginosa, Mol Microbiol 72, 183–201.

[28] Firoved, A. M., Ornatowski, W., and Deretic, V. (2004) Microarray analysis reveals induction of lipoprotein genes in mucoid Pseudomonas aeruginosa: implications for inflammation in cystic fibrosis, Infect Immun 72, 5012–5018.

[29] Sultan, M., Arya, R., and Kim, K. K. (2021) Roles of Two-Component Systems in Pseudomonas aeruginosa Virulence, Int J Mol Sci 22.

[30] Kawalek, A., Glabski, K., Bartosik, A. A., Wozniak, D., Kusiak, M., Gawor, J., Zuchniewicz, K., and Jagura-Burdzy, G. (2023) Diverse Partners of the Partitioning ParB Protein in Pseudomonas aeruginosa, Microbiol Spectr 11, e0428922.

[31] Bignell, C., and Thomas, C. M. (2001) The bacterial ParA-ParB partitioning proteins, J Biotechnol 91, 1–34.

[32] Mishra, D., and Srinivasan, R. (2022) Catching a Walker in the Act-DNA Partitioning by ParA Family of Proteins, Front Microbiol 13, 856547.

[33] O’Neal, L., Baraquet, C., Suo, Z., Dreifus, J. E., Peng, Y., Raivio, T. L., Wozniak, D. J., Harwood, C. S., and Parsek, M. R. (2022) The Wsp system of Pseudomonas aeruginosa links surface sensing and cell envelope stress, Proc Natl Acad Sci U S A 119, e2117633119.

[34] Hogardt, M., and Heesemann, J. (2010) Adaptation of Pseudomonas aeruginosa during persistence in the cystic fibrosis lung, pp 557–562.

[35] Amiel, E., Lovewell Rustin, R., O’Toole George, A., Hogan Deborah, A., and Berwin, B. (2010) Pseudomonas aeruginosa evasion of phagocytosis is mediated by loss of swimming motility and is independent of flagellum expression, Infection and Immunity 78, 2937--2945.

[36] Ballok, A. E., Bahl, C. D., Dolben, E. L., Lindsay, A. K., St Laurent, J. D., Hogan, D. A., Madden, D. R., and O’Toole, G. A. (2012) Epoxide-mediated CifR repression of cif gene expression utilizes two binding sites in Pseudomonas aeruginosa, J Bacteriol 194, 5315–5324.

[37] Winsor, G. L., Griffiths, E. J., Lo, R., Dhillon, B. K., Shay, J. A., and Brinkman, F. S. (2016) Enhanced annotations and features for comparing thousands of Pseudomonas genomes in the Pseudomonas genome database, Nucleic Acids Res 44, D646–653.

[38] Szklarczyk, D., Kirsch, R., Koutrouli, M., Nastou, K., Mehryary, F., Hachilif, R., Gable, A. L., Fang, T., Doncheva, N. T., Pyysalo, S., Bork, P., Jensen, L. J., and von Mering, C. (2023) The STRING database in 2023: protein-protein association networks and functional enrichment analyses for any sequenced genome of interest, Nucleic Acids Res 51, D638–D646.

[39] Ranjani, J., Sivakumar, R., Gunasekaran, P., Velmurugan, G., Ramasamy, S., and Rajendhran, J. (2022) Genome-wide identification of genetic requirements of Pseudomonas aeruginosa PAO1 for rat cardiomyocyte (H9C2) infection by insertion sequencing, Infect Genet Evol 98, 105231.

[40] Zaoui, C., Overhage, J., Lons, D., Zimmermann, A., Musken, M., Bielecki, P., Pustelny, C., Becker, T., Nimtz, M., and Haussler, S. (2012) An orphan sensor kinase controls quinolone signal production via MexT in Pseudomonas aeruginosa, Mol Microbiol 83, 536–547.

[41] Stover, C. K., Pham, X. Q., Erwin, A. L., Mizoguchi, S. D., Warrener, P., Hickey, M. J., Brinkman, F. S., Hufnagle, W. O., Kowalik, D. J., Lagrou, M., Garber, R. L., Goltry, L., Tolentino, E., Westbrock-Wadman, S., Yuan, Y., Brody, L. L., Coulter, S. N., Folger, K. R., Kas, A., Larbig, K., Lim, R., Smith, K., Spencer, D., Wong, G. K., Wu, Z., Paulsen, I. T., Reizer, J., Saier, M. H., Hancock, R. E., Lory, S., and Olson, M. V. (2000) Complete genome sequence of Pseudomonas aeruginosa PAO1, an opportunistic pathogen, Nature 406, 959–964.

[42] Hoboth, C., Hoffmann, R., Eichner, A., Henke, C., Schmoldt, S., Imhof, A., Heesemann, J., and Hogardt, M. (2009) Dynamics of adaptive microevolution of hypermutable Pseudomonas aeruginosa during chronic pulmonary infection in patients with cystic fibrosis, J Infect Dis 200, 118–130.

[43] Sriramulu, D. D., Nimtz, M., and Romling, U. (2005) Proteome analysis reveals adaptation of Pseudomonas aeruginosa to the cystic fibrosis lung environment, Proteomics 5, 3712–3721.

[44] Jacobs, M. A., Alwood, A., Thaipisuttikul, I., Spencer, D., Haugen, E., Ernst, S., Will, O., Kaul, R., Raymond, C., Levy, R., Chun-Rong, L., Guenthner, D., Bovee, D., Olson, M. V., and Manoil, C. (2003) Comprehensive transposon mutant library of Pseudomonas aeruginosa, Proc Natl Acad Sci U S A 100, 14339–14344.

[45] Lee, S. A., Gallagher, L. A., Thongdee, M., Staudinger, B. J., Lippman, S., Singh, P. K., and Manoil, C. (2015) General and condition-specific essential functions of Pseudomonas aeruginosa, Proc Natl Acad Sci U S A 112, 5189–5194.

